# Dbx1 controls the development of astrocytes of the intermediate spinal cord by modulating Notch signaling

**DOI:** 10.1101/2022.03.13.484175

**Authors:** M. Micaela Sartoretti, Carla A. Campetella, Guillermo M. Lanuza

## Abstract

Significant progress has been made in elucidating the basic principles that govern neuronal specification in the developing central nervous system. In contrast, much less is known about the origin of astrocytic diversity. Here we demonstrate that a restricted pool of progenitors in the mouse spinal cord, expressing the transcription factor Dbx1, produces a subset of astrocytes, in addition to interneurons. Ventral p0-derived astrocytes (vA0) exclusively populate intermediate regions of spinal cord with extraordinary precision. Postnatal vA0 population comprises gray matter protoplasmic and white matter fibrous astrocytes and a group of cells with strict radial morphology contacting the pia. We identified that vA0 cells in the lateral funiculus are distinguished by the expression of Reelin and Kcnmb4. We show that Dbx1 mutants have increased vA0 cells at the expense of p0-derived interneurons. Manipulation of the Notch pathway, together with the alteration in their ligands seen in Dbx1 knock-outs, suggest that Dbx1 controls neuron-glial balance by modulating Notch-dependent cell interactions. In summary, this study highlights that restricted progenitors in dorsal-ventral neural tube produce region-specific astrocytic subgroups and that progenitor transcriptional programs highly influence glial fate and are instrumental in creating astrocyte diversity.

## Introduction

Astrocytes, oligodendrocytes and ependymocytes represent about 60% and 75% of spinal cord cells in rodents and humans, respectively (Bjugn and Gundersen, 1993; Fu et al., 2013; Bahney and von Bartheld, 2018). Astrocytes play crucial supportive and active roles in the organization and functioning of the nervous system. They maintain the blood-brain barrier, the distribution of ions and protons, and regulate the water balance. They are also key in the formation, function and plasticity of neural networks by adjusting calcium fluxes, providing neurotransmitter precursors, producing neuropeptides and trophic support factors, and modulating synaptic assembly and transmission (Allen and Eroglu, 2017; Barres, 2008; Verkhratsky and Nedergaard, 2018).

During neural tube development, glial cells are produced from ventricular zone (vz) progenitors that previously generated neurons (Rowitch and Kriegstein, 2010). Substantial advances have been reached in understanding the gene-regulatory programs that control neurogenesis and neuronal diversity in the embryonic spinal cord. It is well established that morphogens pattern the dorsal-ventral (DV) axis of the neural tube, and induce the expression of a combination of transcription factors in spatially limited territories subdividing the neuroepithelium into eleven progenitor domains (Briscoe et al., 2000; Jessell, 2000; Balaskas et al., 2012; Lek et al., 2010; Sagner and Briscoe, 2019). These DV restricted progenitors, p0-p3, pMN and dp1-6, produce distinct cardinal classes of spinal neurons, ventral V0-V3, motoneurons, and dorsal dI1-dI6 and dILA/B (Lai et al., 2016; Lu et al., 2015; Sagner and Briscoe, 2019).

After neurogenesis, the remaining progenitor cells in the vz start to generate astrocyte and oligodendrocyte precursors (Rowitch and Kriegstein, 2010). Together with this switch in competence, the neuroepithelium undergoes changes in gene expression and a phenotypical transformation into radial glia (Barry and McDermott, 2005; Deneen et al., 2006; Stolt et al., 2003; Kang et al., 2012; Freeman, 2010). Glial cells production also follows DV regional organization. Oligodendrocytes, which populate the entire spinal cord, derive from the ventral pMN/pOL and dorsal domains (Zhou et al., 2000; Lu et al., 2000; Cai et al., 2005; Vallstedt et al., 2005; Fogarty et al., 2005). Astrocytes, however, are produced from vz territories spanning the whole DV axis (Rowitch and Kriegstein, 2010; Pringle et al., 2003).

In contrast with the deep knowledge attained on neuronal cell type specification, the comprehension of the developmental principles behind astrocyte specification and diversity is more limited. Recent studies in the embryonic spinal cord and brain indicate that astrocyte development follows a segmental template similar to that involved in early neuron production (Tsai et al., 2012; Vue et al., 2014; Molofsky et al., 2014; Herrero-Navarro et al., 2021).

In the ventral and dorsal spinal cord, different progenitor domains contribute to astrocyte subtypes, which are allocated to spatial regions in accordance with their embryonic origin (Tsai et al., 2012; Hochstim et al., 2008; Vue et al., 2014). At least three molecularly distinct subtypes of ventral astrocytes (vA1, vA2 and vA3) were identified based on the combinatorial expression of the axon guidance and migration proteins Slit1 and Reelin (Hochstim et al., 2008). The spatial arrangement of these populations within the spinal cord grey (GM) and white matter (WM) correlates to their ventricular source (p1, p2 and p3, respectively) (Hochstim et al., 2008; Tsai et al., 2012). In addition, key transcription factors of the gene regulatory networks that control neuron diversification also contribute to astrocyte specification and heterogeneity in the spinal cord. For instance, the bHLH protein Tal1 is necessary and sufficient for the development of astrocytes and the suppression of oligodendrocyte fate in p2 progenitors (Muroyama et al., 2005). Similarly, deletion and forced-expression of Pax6 and Nkx6.1 disrupt the molecular identity of astrocyte subsets in the ventral and ventro-lateral WM (Hochstim et al., 2008; Zhao et al., 2014). Despite this progress, our understanding of the molecular and functional astrocytic heterogeneity across central nervous system regions is far from complete.

Here we describe the specification, migration and maturation of a spinal cord astrocyte population originating from Dbx1-expressing vz progenitors of the p0 domain. Through cell fate tracings, we show that ventral p0-derived astrocytes, vA0, are settled in remarkably precise and reproducible positions to cover the intermediate portion of the mouse spinal cord. This cell group is morphologically heterogeneous and comprises protoplasmic-like astrocytes in the GM, fibrous WM astrocytes and radial astrocytes at the subpial surface.

We further show that in the absence of Dbx1, p0 cells give rise to a significantly increased number of astrocytes, which are produced at the expense of V0 interneurons. We provide evidence that Dbx1 controls the appropriate size of the p0-derived populations by modulating Notch signaling efficacy within the p0 domain and thus coordinating region-specific cell fate developmental features.

## Results

### Dbx1+ progenitors produce glial cells that populate the medial-lateral spinal cord

The transcription factor Dbx1 is expressed in a discrete domain of the neural tube (Fig.1A). To determine the complete contribution of Dbx1-expressing progenitors to spinal cord cell types, we used *Dbx1*^*lacZ*^ mice, where nuclear β-gal recapitulates Dbx1 expression and its stability allows for accurate fate mapping (Fig.1A,B, Pierani et al., 2001; Lanuza et al., 2004; Bouvier et al., 2010). At E13.5, the end of the neurogenic period, V0 interneurons are established in the ventromedial spinal cord (Fig.1C). Remarkably, at E18.5 a different cohort of β-gal^+^ cells is seen in the intermediate region of the spinal cord (Fig.1D arrow, S1D). This group shows a thinner nuclear shape (Fig.1 E,F, Fig S1 A-C) and a specific expression profile. They are negative for neuronal NeuN and express Sox2 (Fig.1G-J), a transcription factor active in progenitors and glial cells (Hoffmann et al., 2014).

**Figure 1.**
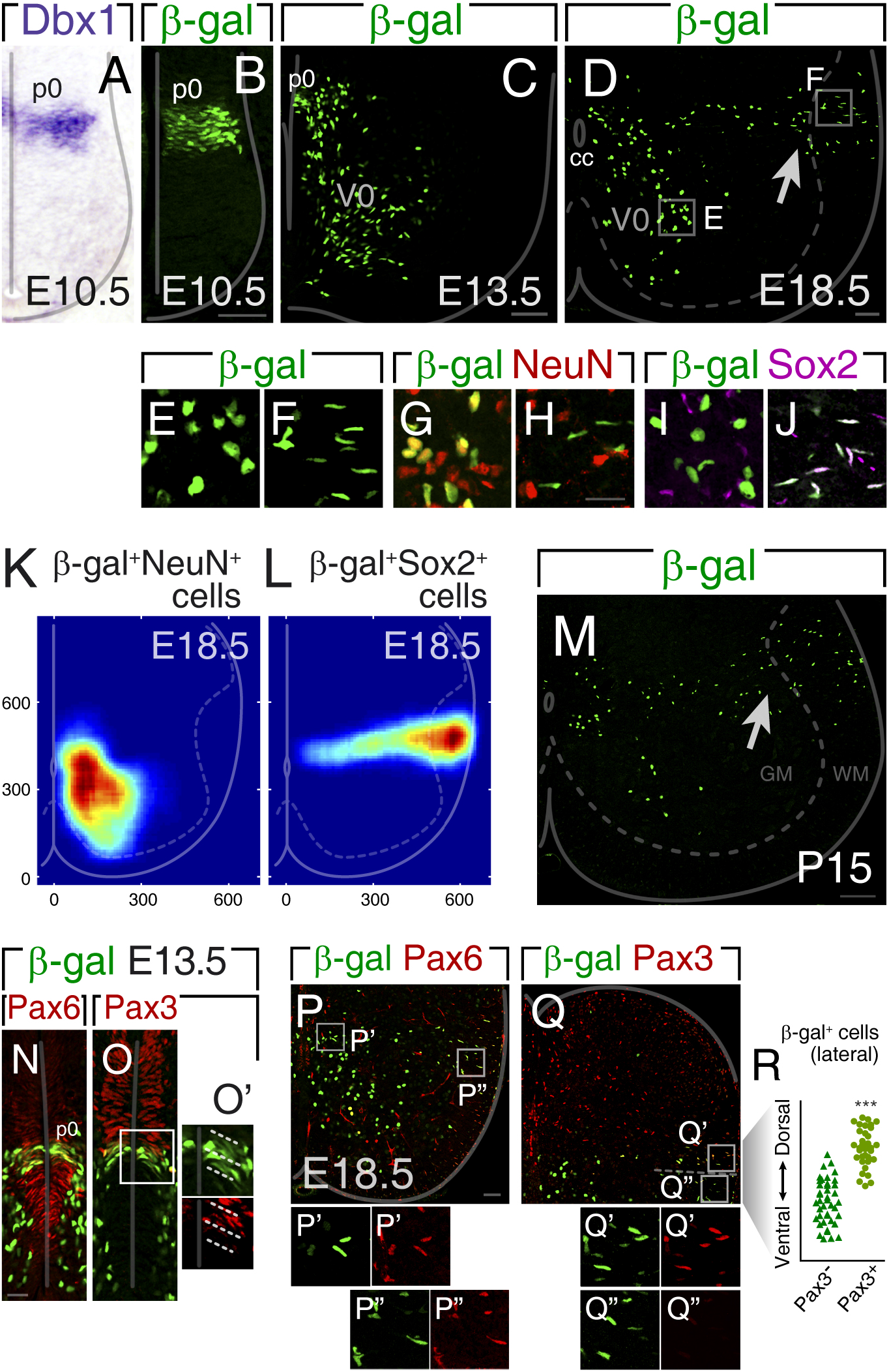
Dbx1+ p0 progenitors produce glial cells that populate a defined region of the intermediate-lateral spinal cord. (A-J) Dbx1-derived cells in *Dbx1lacZ* mice. (A,B) Transverse E10.5 neural tube sections hybridized with Dbx1 probe and immunostained against β-gal. (C) At E13.5 β-gal+ population includes V0 interneurons and late progenitors in the p0 domain. (D-J) E18.5 cross-sections showing V0 neurons and a group of β-gal+ cells between the midline and the *lateral funiculus* (arrow). (E-J) Dbx1-derived neurons (E,G,I) and β-gal+ lateral cells (F,H,J) have distinctive nuclei morphology, NeuN and Sox2 expression. (K-L) Density maps of β-gal+ cells in E18.5 spinal cord based on the location of 890 NeuN+ and 775 Sox2+ cells. (M) The regional allocation of β-gal+ astroglial cells is maintained postnatally as seen in P15 spinal cord. (N-O’) E13.5 p0 vz progenitors are Pax6+ and are subdivided in dorsal and ventral pools based on Pax3 expression. (P-R) DV organization of glial populations at E18.5. β-gal+ cells are Pax6+. β-gal+ cells positioned more dorsally express Pax3 (Q’,R) while cells settled ventrally are negative (Q”,R) (***p<0,001, Mann Whitney test). Scale bars: 50μm in A-D,P,Q; 20μm in E-J,N,O; 100μm in M. Data: mean±SD. See also Figure S1.

Quantitative distribution analyses of β-gal^+^,Sox2^+^ cells revealed that p0-derived glia colonize a precise region of the spinal cord (Fig.1L, S1 F). β-gal^+^,Sox2^+^ nuclei stand between the midline and the pial surface, spanning the GM and WM (Fig.1L, S1G-J). This territory is different from V0 location (Fig.1K, S1E), indicating that neurons and glia produced by the same vz domain follow dissimilar migration routes. In the hindbrain, p0 cells also generate both neurons and glia, although intermingled in the ventrolateral medulla (Fig.S1K-N).

Lateral β-gal^+^ cells are only seen at advanced embryonic stages, suggesting they are produced after E13, once neurogenesis has ended. At E13.5, vz Dbx1 spatial expression (Fig.1N,O, Fig.S1O) remains similar to its earlier pattern (Briscoe et al., 2000; Pierani et al., 2001). Stainings for transcription factors with DV restriction show vz β-gal^+^ cells within Pax6^+^ territories, dorsal to Nkx6.1^+^ domains (Fig.1N, Fig.S1O). The dorsal patterning gene Pax3 partially overlaps with Dbx1, establishing ventral and dorsal p0 subdomains (Fig.1O,O’). At E18.5, β-gal^+^ glial cells in the GM and WM are Pax6^+^ (Fig.1P-P”, ~96%) and do not express Nkx6.1, which marks more ventral WM cells (Fig.S1P-P”, Hochstim et al., 2008). We also found that 43±4% of Dbx1 progeny is Pax3^+^ (Fig.1Q-Q”), being Pax3 limited to more dorsal β-gal^+^ cells (Fig 1 Q-R). The uneven Pax3 expression strictly correlates with the polarized early Pax3 patterning and indicates that the arrangement of β-gal^+^ glia closely mirrors their DV vz origins. To determine if the precise positioning of Dbx1-derived glia is a transient state to later disperse in the spinal tissue, we analyzed their postnatal location. However, β-gal^+^ cells at P6 and P15 are still restricted to the intermediate region without DV displacement (Fig.1M, S1Q).

In summary, fate mappings indicate that Dbx1^+^ progenitors produce, in addition to V0 neurons, a population of glial cells that migrate laterally and settle in a specific region of the postnatal spinal cord.

### Dbx1-derived population comprises protoplasmic, fibrous and radial astrocytes

To explore the identity of p0-derived cells, we studied the expression of several molecular markers. In newborn pups, all β-gal^+^,Sox2^+^ cells robustly co-labeled with the astroglial transcription factors Sox9 and Nfia (Deneen et al., 2006; Kang et al., 2012) (Fig.1J, Fig.2A-C,H). The glutamate transporter Glast, restricted to the vz and the astrocytic lineage (Shibata et al., 1997), was expressed in most β-gal^+^ cells at P0, in line with their astroglial identity (Fig.2D,H). As shown above, NeuN was not detected in β-gal^+^ cells in the intermediate-lateral cord (Fig.1H, Fig.2E,H). Since previous studies have established oligodendrocyte production from dorsal domains (Cai et al., 2005; Fogarty et al., 2005; Vallstedt et al., 2005), we examined if some β-gal^+^ cells were oligodendrocyte precursor cells. However, we found they were Olig2 negative (Fig.2F,H). To fully confirm the identity of p0-derived cells, we assessed the presence of the astrocytic protein Gfap. We found that β-gal^+^ cells within or close to the WM displayed Gfap^High^ processes, while those in the GM had very low signal (Fig.2G,H). The expression of these proteins was also analyzed at P6 and P15, confirming their astrocytic nature. Similar to P0, at p6 and P15, β-gal co-labeled with Sox9, Sox2 and Nfia (Fig.S2 A-C,E), while remaining NeuN- and Olig2-negative (Fig.S2 D,G,H). At P15, β-gal^+^ cells were Gfap-positive, with high expression in WM cells (Fig.S2 F). Thus, marker analysis indicates that p0 progenitors produce cells that differentiate into astrocytes (vA0).

**Figure 2.**
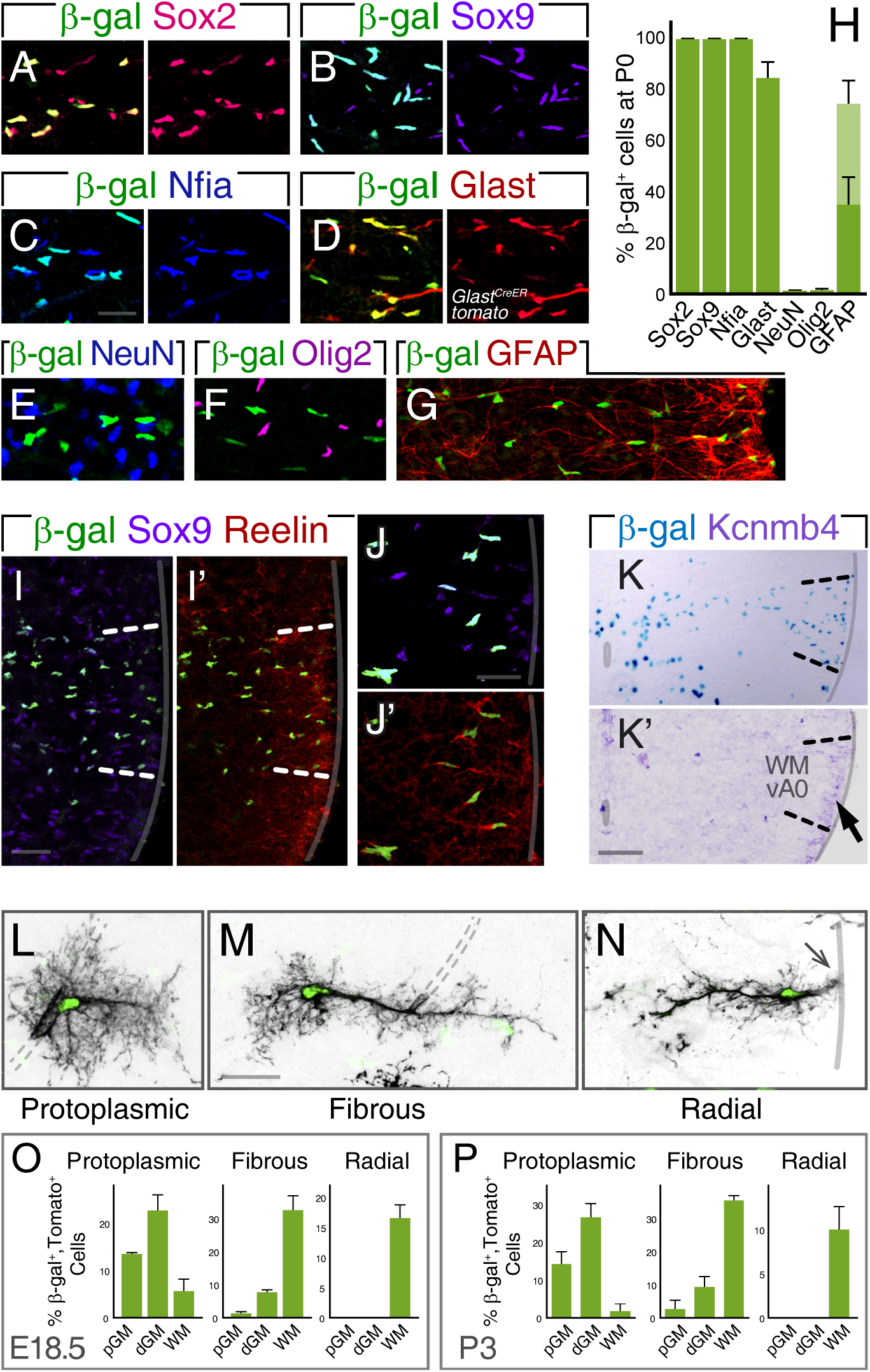
p0-derived vA0 population is composed by protoplasmic, fibrous and radial astrocytes. (A-H) β-gal cells in the intermediate spinal cord express astrocytic markers. (A-G) P0 spinal cord stained against β-gal, Sox2, Sox9, Nfia, Glast (*GlastCreER*;*CAG:LSL-tdTomato*, Tam @E18.5), NeuN, Olig2 or GFAP. (H) Percentage of β-gal+ cells expressing each protein (12 sections, 3 cords). (I-K) vA0 cells in the WM express Reelin and Kcnmb4. (I,J) P0 cross-sections stained for β-gal, Reelin and Sox9. (K) *In situ* hibridization for Kcnmb4 showing expression in vA0 area close to the pia. (L-P) vA0 population comprises protoplasmic, fibrous and radial astrocytes. (L-N) Representative examples of Dbx1-derived astroglia at P3. Cells were identified by nuclear β-gal and tomato using *Nestin:CreER;CAG:LSL-tdTomato* mice induced with low dose of Tam at E11.75. Dotted lines demarcate blood vessels. (O,P) Percentages of β-gal+ cells along the medial-lateral axis (pGM, dGM, WM) at E18.5 and P3 (296 cells from 11 E18.5 embryos and 92 cells from 3 P3 pups). Scale bars: 20μm except 100μm in K. Values are mean±SD. See also Figure S2.

The glycoprotein Reelin is co-expressed with Pax6 in vA1 and vA2 WM astrocytes (Hochstim et al., 2008). As vA0 cells are also Pax6^+^ (Fig.1P), we analyzed its presence in vA0 population, and found that Reelin labels WM Dbx1-derived cells (Fig.2I-J’, Fig.S2I-L).

In search of novel vA0 markers, a survey of the Allen Spinal Cord Atlas (http://mousespinal.brain-map.org/) pointed at Kcnmb4, the auxiliary β4-subunit of BK channels (Contet et al., 2016), expressed in the lateral funiculus. *In situ* hybridizations showed Kcnmb4 expression in the WM occupied by vA0s (Fig.2K). Similar to Reelin, Kcnmb4 is limited to WM marginal cells (Fig.2K-K’). At E13.5, Reelin is absent in vz Dbx1^+^ cells (Fig.S2M), while Kcnmb4 is already expressed in the p0 domain (Fig.S2N,O). Hence, Reelin and Kcnm4 are distinctive molecular markers of vA0 population.

Since vA0 cells are distributed throughout the GM and WM, we evaluated their morphology to determine whether Dbx1-derived population includes the major classes of astrocytes: protoplasmic and fibrous (Oberheim et al., 2012; Peters et al., 1991). For this purpose, we generated animals carrying the transgene *Nestin:CreER* and the conditional tdTomato reporter and induced scattered recombination in the vz to identify later isolated cells in the parenchyma (Fig.S2P-Q”). There were three major vA0 morphologies at E18.5 and P3. First, 42-43% of β-gal^+^ cells were protoplasmic-like astrocytes with highly branched bushy processes emanating from their soma (Fig.2L). Second, 42-47% of vA0 cells were fibrous-like astrocytes with a main straight long radial process and shorter lateral ramifications (Fig.2M). We found that both classes have cellular extensions surrounding blood vessels (Fig.2L,M dotted lines, Tabata, 2015). Finally, 10-16% of β-gal^+^ cells, whose nuclei localized at the subpial surface, display a pronounced radial morphology with short transverse processes and contacting the pia (Fig.2N, arrow) (Liuzzi and Miller, 1987; Petit et al., 2011; Barry and McDermott, 2005).

To define regions occupied by different vA0 subtypes, they were classified according to their medial-lateral location into proximal or distal GM (pGM, dGM) and WM. Consistently, the majority of protoplasmic-like vA0s were in the pGM and dGM (Fig.2O,P, ~90%), while fibrous-like were mainly in the WM (Fig.2O,P, ~80%). Finally, β-gal^+^ astrocytes with strict radial shape were only settled in the WM close to the pia (Fig.2O,P). These results demonstrate that Dbx1 progenitors produce a heterogeneous population of astrocytes that occupy defined regions of the spinal cord GM and WM.

### vA0 precursors derive from radial glia and progressively occupy lateral spinal regions

Experiments above suggest that astroglia arises from Dbx1^+^ vz cells after E13.5. Spinal cord neuroepithelial progenitors acquire the radial glia phenotype before becoming the source of glial lineages (Barry and McDermott, 2005). Immunostaining against the radial glia marker Nestin, shows that E13.5 β-gal^+^ ventricular cells radiate long basal processes reaching the basal lamina (Fig.3A). Additionally, β-gal^+^ vz cells co-express Sox2, Sox9 and Blbp, another radial glia marker (Barry and McDermott, 2005) (Fig.3B-D). To confirm that vA0 population derives from radial glia, we analyzed the fate of Dbx1^+^,Glast^+^ cells using *Glast^CreER^*;*tdTomato* conditional reporter mice. Glast appears in spinal cord radial glia slightly preceding the gliogenic period (Shibata et al., 1997). Indelible marking, induced at E13.5, activated recombination in ~90% of germinal zone cells but not in the mantle (Fig.S3 A-C). As predicted, at E18.5 we found that β-gal^+^ cells that migrated laterally were Tomato^+^ (~90%, Fig.3E-E’), demonstrating that Glast^+^ radial glial cells in the E13.5 p0 domain are the source of vA0 astrocytes.

**Figure 3.**
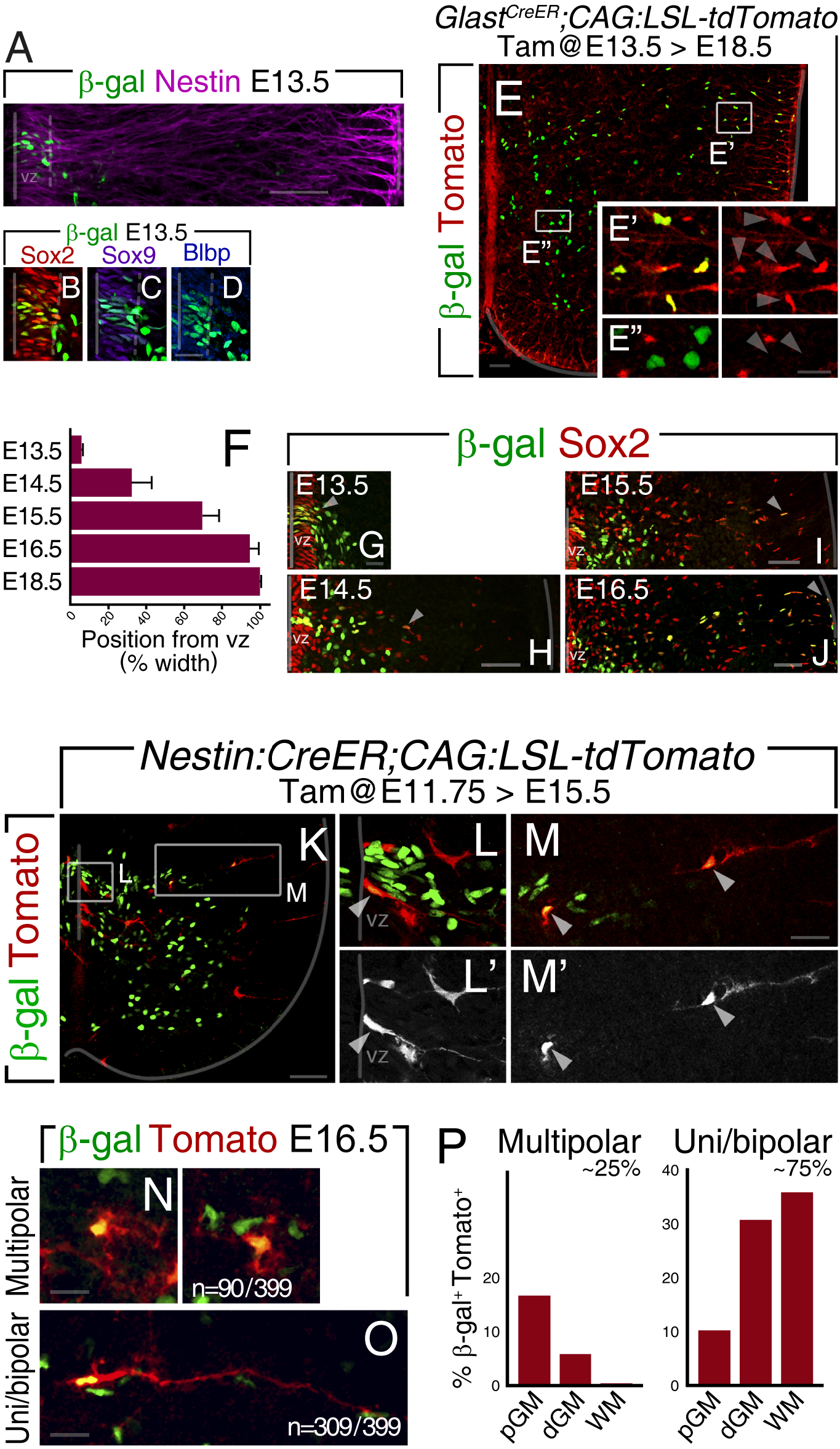
Dbx1 radial glia gives rise to astrocyte precursors that progressively occupy lateral spinal regions. (A-E”) Delayed p0 lineage derives from spinal cord radial glia. (A-D) β-gal, Nestin, Sox2, Sox9 and Blbp staining at E13.5. (E-E”) Late-born β-gal+ cells are produced from E13.5 Glast+ progenitors. Tomato labeling in *GlastCreER;CAG:LSL-tdTomato* embryos was induced with Tam at E13.5 and analyzed at E18.5. β-gal+ cells in E18.5 lateral spinal cord are tomato+ (89±6%, E’) (653cells, 9 sections, 3 embryos), while V0 neurons are negative (E”). (F-J) Dbx1-derived astroglia progressively occupies distal spinal cord regions. (F) Relative distance of front runner β-gal+,Sox2+ nuclei (top 10%) from the vz border. (G-J) Staining of E13.5-E16.5 spinal cords for β-gal and Sox2. Arrowheads point out the most distal cells. (K-P) Developing Dbx1-derived astroglial cells are morphologically heterogeneous. Mosaic tomato labeling was induced by low Tam doses at E11.75 in *Nestin:CreER;CAG:LSL-tdTomato* and analyzed at E15.5 and E16.5. (K-M) E15.5 spinal section stained for Tomato and β-gal. Magnification of p0 domain with a β-gal+,Tomato+ cell in the vz (arrowhead, L) and two cells with dissimilar morphologies in the mantle zone (arrowheads, M). (N,O) Representative images of cell morphologies at E16.5. Multipolar cells accounted for 23% (n=90/399), while cells with unipolar or bipolar morphology were 77% (n=309/399). (P) Multipolar cells were mainly in the pGM, while uni/bipolar β-gal cells were more lateral, mostly in dGM and WM. Scale bars: 50μm in A,E,H-J,K; 20μm in B-D, E’,E”,G,L-M’; 10μm in N,O. Values are mean±SD.

We next characterize vA0 development in more detail. At E13.5 only a few β-gal^+^,Sox2^+^ cells were spotted just outside the vz (Fig.3G). Later, they gradually occupied more distal positions, reaching the pial surface by E16.5 (Fig.3F-J). Remarkably, β-gal^+^,Sox2^+^ nuclei were restricted to DV intermediate coordinates during development, suggesting exclusive radial migration. At E14.5 and E16.5, β-gal^+^,Sox2^+^ cells moving laterally also express Sox9 and Nfia (Fig.S3 D-H).

To evaluate how Dbx1-derived cells migrate, we studied their shape by mosaic labeling. At E15.5, some β-gal^+^ cells with thin radial process were still in the vz (Fig.3K,L). In the mantle zone, two contrasting morphologies were seen: cells with short processes and others with prolonged extensions toward the pia (Fig.3M, arrowhead). Further analysis at E16.5, when p0-progeny already spans the whole medial-lateral axis, reinforced the presence of two main shapes. First, multipolar cells displaying short ramifications (Fig.3N) accounted for ~25% and were mainly in the pGM (Fig.3P). Second, uni/bipolar cells with one extension towards the pia with or without an apically-directed process (Fig.3O) represented ~75% and were predominantly in the dGM and WM (Fig.3P). The morphology of uni/bipolar cells contacting the pia, together with their soma displacement, are reminiscent of radial migration processes previously described (Nadarajah et al., 2001; Pakan and McDermott, 2014).

### vA0 precursors intensively proliferate distal from the vz

Quantification of β-gal^+^ cells in the parenchyma revealed a continued increase from E13.5 to E18.5 (Fig.4A). This rise was accompanied by attrition of the p0 vz, which at E16.5 contains few remaining cells (Fig.4B). To determine the expansion of vA0 population during the migratory journey, we assessed their proliferation with BrdU. At E13.5, about 7% of β-gal^+^ vz cells incorporated BrdU (Fig.4C). Later, β-gal^+^,BrdU^+^ cells drastically increased in the mantle zone reaching a proliferation peak at E16.5 (24.1±8.5%, Fig.4C,D). These stages with high division rates in the mantle coincide with the largest expansion of vA0 cells (Fig.4A). Proliferation at E16.5 was not restricted to any preferential morphology, with multipolar and uni/bipolar cells incorporating BrdU (~23-25%, Fig.S3I-K). Cell divisions were largely biased to distal positions, as 90% of BrdU-labeled cells were in the dGM and WM (Fig.4E-G). Further analysis revealed that proliferation persists at E18.5 although significantly decreased (E18.5: 7.8±2.1%, Fig.4C) and β-gal^+^,BrdU^+^ cells were mostly in the WM (>80%, Fig.4H,I). In P2, P6 and P15 animals, BrdU^+^ cells were very reduced (Fig.4C and not shown). Overall, these experiments show that vz Dbx1^+^ radial glia produces heterogeneous intermediate astrocyte precursors, which intensively amplify outside the germinal zone in distal regions of the spinal cord to shape the postnatal vA0 population.

**Figure 4.**
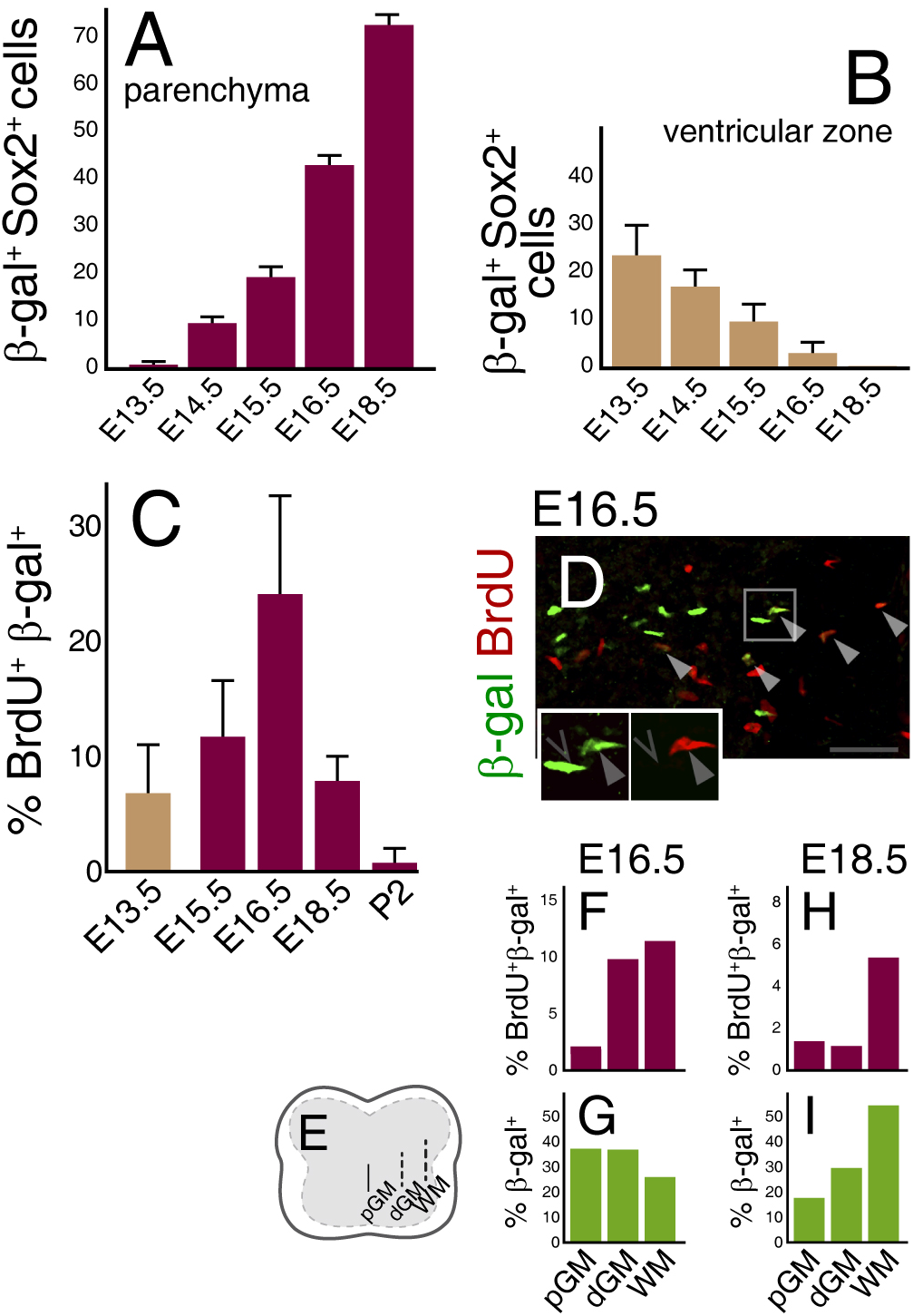
vA0 cells extensively divide during migration. (A-B) Quantification of β-gal+,Sox2+ cells in the E13.5-E18.5 spinal cord mantle zone (A) or within the vz (B); (>20 sections,>3 embryos). (C-I) β-gal+ population expands outside the vz. (C) Percentage of β-gal+ cells that incorporated BrdU at different stages (>10 sections, 2-3 embryos). (D) E16.5 spinal cord with β-gal and BrdU; colocalizations. (E-I) Proportion of β-gal+,BrdU+ cells in medial-lateral regions (F,H) and distribution of β-gal+ cells (G,I) at E16.5 and E18.5 (n=377 and 615 cells, respectively). Scale bars: 50μm. Data: mean±SD. See also Figure S3.

### Dbx1 regulates the development of vA0 population

Dbx1 plays important functions in the specification of V0 neurons in the spinal cord and hindbrain (Pierani et al., 2001; Lanuza et al., 2004; Zagoraiou et al., 2009; Bouvier et al., 2010). To evaluate the role of Dbx1 in the differentiation of the late p0 progeny, we produced *Dbx1 mutants* in which the *Dbx1*^*lacZ*^ reporter/null allele allows cell tracing. We first obtained E18.5 *Dbx1* heterozygotes and mutants and found a significant expansion of β-gal^+^,Sox9^+^ cell number in the *Dbx1-KO* spinal cord (Fig.5A-C; 35±13% increase). To better characterize the mutant vA0 population, we studied its spatial distribution and saw that *Dbx1*^−/−^ β-gal^+^ cells were confined to a region similar to *controls* (Fig.5A,B,E,F, Fig.S4A). In addition, their radial positions from the midline to the lateral funiculus were unchanged (Fig.5E-G). We spotted a slight ventral angular shift (Fig.S4A,B) which is likely consequence of lamina VIII reduction in *Dbx1-KOs*. As in *controls*, β-gal^+^ cells in *Dbx1-KOs* express Sox2, Sox9 and Nfia (Fig.5H, Fig.S4 H-J) being Olig2- and NeuN-negative (Fig.S4H,J). Hence in the absence of Dbx1, late-born p0 cells still possess astrocytic character.

**Figure 5.**
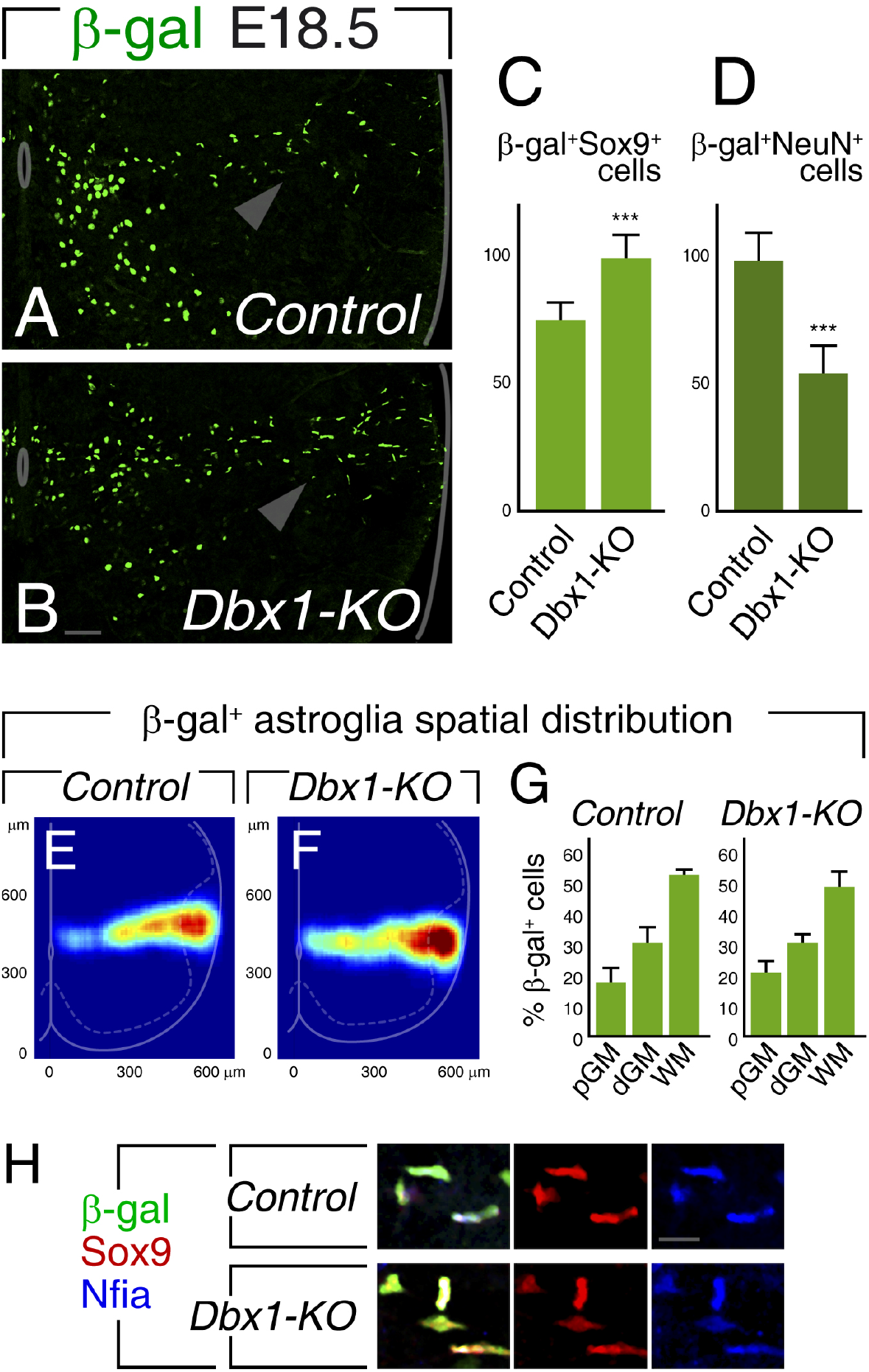
Dbx1 controls the size of astroglial vA0 population. (A-D) *Dbx1* mutants have increased p0-derived astrocytic cells at the expense of V0 neurons. E18.5 *Control* and *Dbx1-KO* spinal cord cross-sections (A-B). Quantification of astroglial β-gal+,Sox9+ (C) and neuronal β-gal+,NeuN+ (D) cells per section, 35±13% increment and 44±9% decline, respectively (n=22 sections from 3 embryos, ***p< 0.001, Mann-Whitney test). (E-F) Spatial maps of β-gal+,Sox9+ cells in *control* and *Dbx1* mutants made from 650 and 895 β-gal+ cell, respectively. (G) The distribution of cells in the medial-lateral axis is similar between genotypes (non-significant, Mann-Whitney test). (H) β-gal+ cells in *control* and *Dbx1−/−* are colabeled with glial markers Sox2 and Nfia. Scale bars: 50μm in A,B; 10μm in H. Values are mean±SD. See also Figure S4.

Interestingly, we noticed that the increment of glial β-gal^+^ cells in *Dbx1-KOs* was accompanied by a decrease in β-gal^+^ neurons in the ventromedial cord (Fig.5A-D). We also found this phenotype in the hindbrain, where Dbx1 mutants had increased β-gal^+^,Sox9^+^ cells, while neurons were reduced (Fig.S4 C-G).

In summary, *Dbx1* mutant spinal cord and hindbrain have an enlarged vA0 population and a diminished number of neurons produced by the same ventricular domain. These results demonstrate that in addition to the specification of V0 interneurons, Dbx1 controls the differentiation of p0-derived astroglial cells.

### Dbx1 defines the prospective astrocytic progenitor pool

The observation that vA0 astrocytes are increased in *Dbx1*^−/−^ prompted us to define when and how Dbx1 acts on astrogliogenesis. Several possibilities could explain this phenotype in *Dbx1-KO*: developing vA0 amplify at higher rates, glial cells are prematurely born, or ventricular p0 precursors are already increased in mutants at the neurogenesis-gliogenesis switch.

First, we analyzed the expression of Dbx1 throughout development and confirmed its expression in the vz and absence in the mantle zone (Fig.6A-E), suggesting Dbx1 acts in progenitors. Second, we quantified the number of β-gal^+^,Sox9^+^ glial cells in the mantle zone of *control* and *mutants* from E13.5 to E18.5. vA0 population was higher in *Dbx1*^−/−^, being their increase already significant at E14.5 (Fig.6F). To define whether the increment rate of developing vA0 during this period is different in *Dbx1*^−/−^, we determined the ratio of β-gal^+^ cells in *Dbx1-KO* over *control*. This number was rather constant at all stages, indicating proportional increases in *control* and *mutants* (Fig.6G, 1.36±0.08). The proliferation patterns in *Dbx1-KO*s and *controls* were not different (not shown). Lastly, we did not observe differences in β-gal^+^,Sox9^+^ cells outside the vz at E13.5 (Fig.6F), ruling out a premature gliogenesis onset in *Dbx1-KO*.

**Figure 6.**
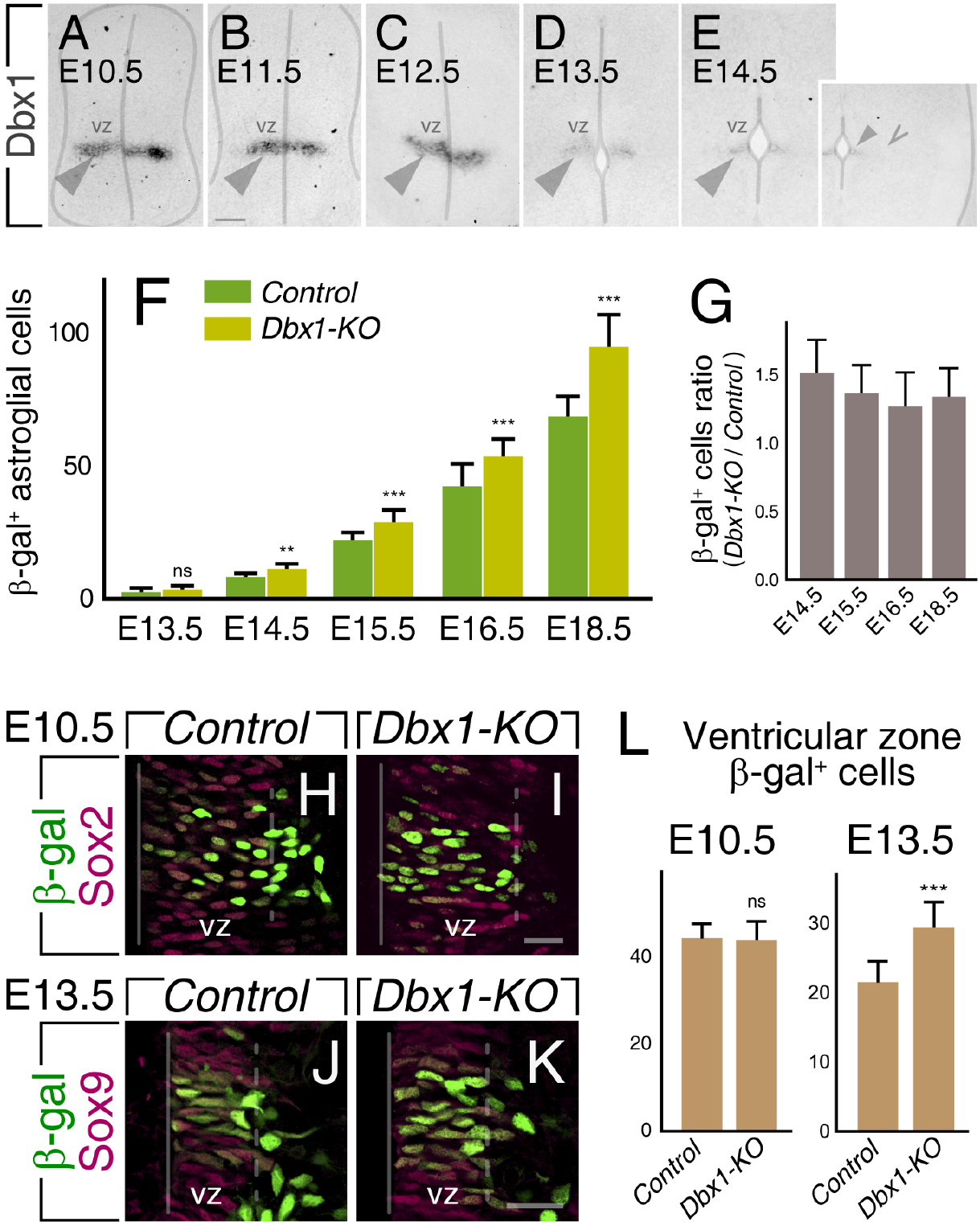
Dbx1 deliniates the p0 progenitor pool and vA0 astroglial population. (A-E) Dbx1 is expressed at p0 vz (filled arrowheads) and absent in the mantle zone (empty arrowhead). (F) Quantification of β-gal+,Sox9+ cells outside the vz in *control* and *Dbx1-null* E13.5-E18.5 cords (>10 sections from 2-3 embryos of each genotype/stage, **p<0.01,***p<0.001, Mann Whitney test). (G) Ratio of β-gal+ glial cells in *Dbx1-KO* to *control* (non-significant, Kruskal-Wallis with *post hoc* Dunn’s test). (H-L) Late p0 vz progenitors are increased in *Dbx1* mutants. (L) Quantification of β-gal+,Sox2/9+ cells in the vz. No differences were seen near the onset of the neurogenic (E10.5), while *Dbx1-KO* have increased β-gal+ progenitors at E13.5 (38±23% increment; 10-16 sections, 2-3 embryos ea.; ns,non-significant, ***p<0.001, Mann Whitney test). Scale bars: 30μm in A-E; 20μm in H-K. Data: mean±SD.

Altogether these results suggest that Dbx1 limits vA0 population by acting early, before p0 start astrocytic production. To test this possibility, we analyzed the vz at the neuron-glia transition. In *Dbx1* mutants, we found β-gal^+^,Sox9^+^ numbers were significantly increased within the E13.5 p0 domain (Fig.6J-L), indicating that the expansion seen in mutant relates to an enlarged late progenitor pool with astroglial potential. To determine when p0 domain increases in *Dbx1*^−/−^, we analyze the E10.5 vz, and found that p0 neuroepithelial progenitor numbers were similar in *Dbx1-KO* and *controls* (Fig.6H,I,L). This indicates that the increase in p0-domain progenitors takes place during the neurogenic phase before gliogenesis begins. Moreover, the proportional increase rate of vA0 cells and the absence of premature vz exit in *Dbx1-KO* eliminates the possibility of Dbx1 controlling vA0 population during the gliogenic phase. We propose that Dbx1-dependent modulation of astrogenesis is a consequence of Dbx1 acting during the neurogenic period, establishing the size of the late progenitor pool with the potential to produce vA0 cells.

### Dbx1 controls astroglial development by directing Notch ligand expression

Changes in neuron-astrocyte balance in *Dbx1* mutants are reminiscent of phenotypes with altered Notch signaling. (Louvi and Artavanis-Tsakonas, 2006; Freeman, 2010). To explore a relation between Dbx1 and Notch signaling in determining late p0 progenitor number and astroglial progeny, we evaluated key components of the Notch-Delta pathway. Previous studies have shown that specific ligand-receptor pairs distinctly activate Notch signaling and that post-translational modifications of Notch are crucial in ligand activation (Hicks et al., 2000; Yang et al., 2005; Kakuda and Haltiwanger, 2017). Notch ligands, Dll1 and Jag1, display a striped expression pattern in DV neural tube domains (Lindsell et al., 1996; Myat et al., 1996; Marklund et al., 2010; Ramos et al., 2010). The p0 domain expresses Dll1, which is also in other ventricular territories (Fig.7A-C,E). Likewise, Jag1 is restricted to the p1 and pd6 domains, bordering dorsally and ventrally the Dbx1^+^ vz (Fig.7A,B,D,F). The glycosyltransferase Lfng, known to modify Notch receptor properties (Hicks et al., 2000; Yang et al., 2005; Kakuda and Haltiwanger, 2017), shows an expression largely similar to Dll1 (Fig.7E,G).

**Figure 7.**
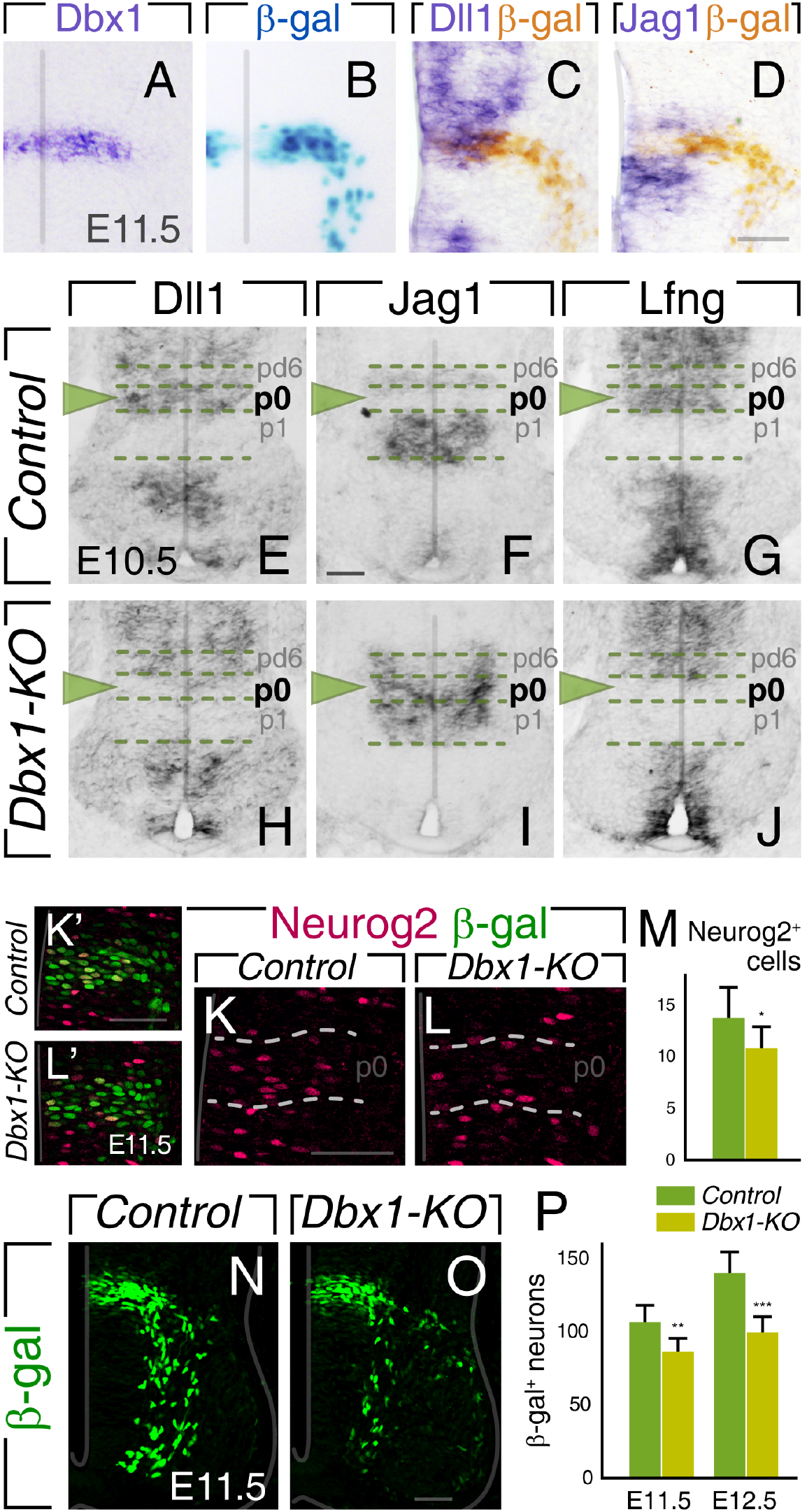
Dbx1 controls the expression of Notch regulatory proteins and neurogenesis in p0 domain. (A-D) Cells within the p0 domain express Dll1 but not Jag1. E11.5 neural tube hybridized for Dbx1, developed for β-gal and hybridized for Dll1 or Jag1 with β-gal immunostaining. (E-J) Dbx1 regulates Notch ligand expression. *In situ* hybridizations at E10.5 show that Dll1 and Lnfg in the p0 region are replaced by Jag1 expression in *Dbx1* mutants. Arrowheads indicate the p0 domain. (K-P) Neurogenesis from p0 progenitors is reduced in *Dbx1-KO*. (K-M) Immunostaining of *control* and *Dbx1-KO* E11.5 spinal cord against Neurog2 and their quantification. (N-P) β-gal+ mantle zone cells in *Dbx1* mutants are diminished at E11.5 and E12.5. Data: mean±SD. Scale bars: 50 μm.

The analysis of Notch ligand patterning in E10.5 *Dbx1-KOs* showed that the p0 region switched to Jag1-positive, Dll1- and Lfng-negative (Fig.7H-J). The altered pattern in *Dbx1* mutants was seen throughout the neurogenic period (E11.5 and E12.5, Fig.S5G-N), while Notch1/2 receptors expression in mutants was unaltered (Fig.S5A-D). Dll3, present in cells fated for terminal neuronal differentiation at the vz-mantle transition (Dunwoodie et al., 1997), is reduced in *Dbx1-KO* likely reflecting diminished p0 neurogenesis (Fig.S5E,F).

These results suggest that in *Dbx1-KOs*, Jag1, expressed in committed neuronal precursors, is more efficient at activating Notch signaling in neighboring progenitors than Dll1 (Hicks et al., 2000). Consistently, we found that *Dbx1* mutants have less Neurog2^+^ neuron-committed cells in their p0 domain (Fig.7K-M), and a reduced population of β-gal^+^ neurons (Fig.7N-P). As shown above, the reduction in the pace of neurogenesis in *Dbx1* mutants is accompanied by an increase in undifferentiated progenitors at the end of the neurogenic period.

### vA0 development is dependent on early Notch signaling

To assess how Notch signaling is involved in regulating V0 and vA0 fates, we generated Presenilin1 (*Psen1*) mutants, in which Notch activation is impaired. At E18.5, the number of vA0 cells in *Psen1*^−/−^ spinal cords is severely reduced (Fig.8A-C, dotted box), while V0 neurons are increased (Fig.8A,B,D), confirming that perturbations in the Notch pathway result in profound alterations in p0 cell identities.

**Figure 8:**
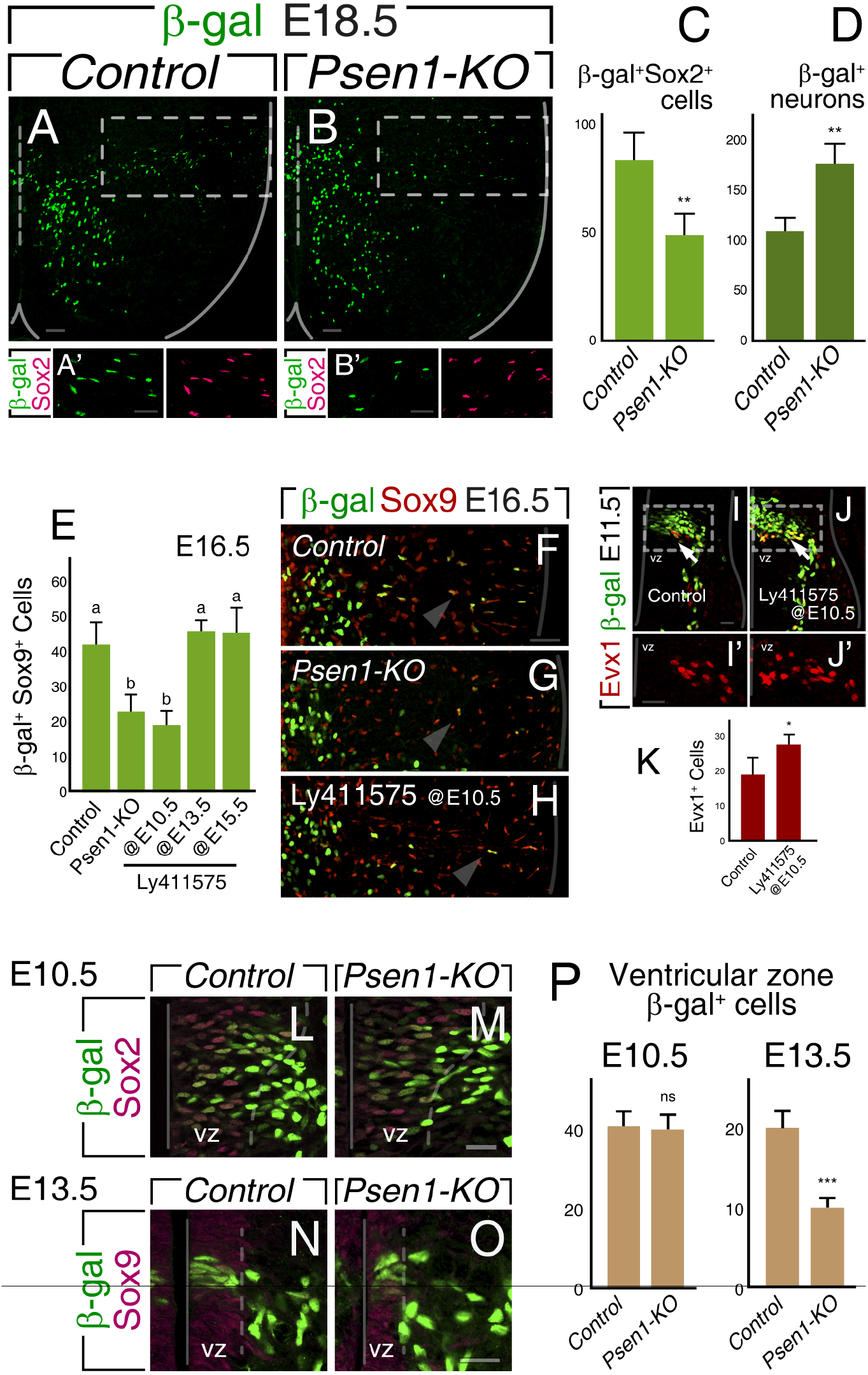
Notch signaling balances glial-neuronal p0 fate. (A-D) vA0 population is decreased after Notch signaling abrogration. E18.5 *Psen1+/+* (*control*) and *Psen1-KO* stained for β-gal and Sox2 (A’,B’). Number of β-gal+,Sox2+ (C) and β-gal+,Sox2-(D, neurons) cells per section; 40±7% reduction and 63±19% increment, respectively (n=12 sections from 3 embryos, **p< 0.01, Mann-Whitney test). (E-K) Early Notch signaling inhibition affects vA0 differentiation. (E-H) Number of β-gal+,Sox9+ cells per section at E16.5 after treating with vehicle (*control*) or Ly411575 at E10.5, E13.5 or E15.5 (n>10 sections, 2-3 embryos). Letters indicate significant differences between groups (p<0.01, Kruskal-Wallis with *post hoc* Dunn’s test). (I-K) Early pharmacological Notch signaling inhibition increases neurogenesis. Evx1 staining in E11.5 embryos treated with vehicle or Ly411575 at E10.5 (*p< 0.05, Mann-Whitney test n=10 sections, 2 embryos ea.). (L-P) Late p0 progenitors are diminished in *Psen1-KO*. Quantification of β-gal+,Sox2/9+ cells in the *control* and *Psen1-KO* vz at E10.5 and E13.5. No difference was found at E10.5, while the p0 cells were significantly decreased at E13.5 (49±6% reduction, 12 sections, 2-3 embryos ea.; ns,non-significant, ***p<0.001, Mann Whitney test). Dotted lines mark vz limits. Scale bars: 50μm in A,B,F-H; 30μm in A’,B’; 20μm in I-J’,L-O. Values are mean±SD. See also Fig. S5.

To define when Notch signaling is relevant in vA0 development, we pharmacologically perturbed Notch processing using the γ-secretase inhibitor Ly411575. Ly411575 was applied at specific stages, at E10.5 (neurogenic phase), E13.5 (neurogenic-gliogenic transition) or E15.5 (astrocytic migration/expansion period). Ly411575 administration at E13.5 or E15.5 did not result in changes in vA0 number (Fig.8E). However, when embryos were treated at E10.5, we found a notable decrease in vA0 cells at E16.5, similar to *Psen1-KOs* (Fig.8E-H). These results confirm that perturbations in Notch signaling in an early developmental window highly influence the specification of p0 astrocyte precursors. To verify that Notch inhibition at E10.5 also modulates neuronal differentiation, we analyzed the V0 marker Evx1 at E11.5. As expected, there was an increase in V0 interneurons, with Evx1-expressing neurons ectopically immersed in the vz, reflecting premature differentiation (Fig.8I-K). Altogether, these experiments demonstrate that early Notch-mediated cell-cell interactions in the p0 domain define not only neuron production, but also late glial development.

We finally examined how abrogation of Notch signaling impacts the p0 pool available for glial differentiation. In *Psen1* mutants, we found a drastic decrease in p0 vz cells at E13.5 (Fig.8N-P). *Psen1-KO*s have normal p0 numbers at E10.5 (Fig.8L,M,P), implying that the differences seen at E13.5 are acquired during the neurogenic phase. These results show that Notch signaling perturbations generate increased and premature neuron production, leading to a reduced late p0 progenitor pool at the beginning of the gliogenic phase. Together with data presented above (Fig.5), these results demonstrate that Dbx1 and Notch modulate p0 progenitor behavior during the neurogenic period, controlling the balance between neuron production and preservation of undifferentiated vz cells with the potential of subsequently give rise to vA0 astrocytes.

## Discussion

This study shows that neural progenitors of the vz p0 domain expressing Dbx1 generate astroglial precursors that cover a specific region of the mouse spinal cord following a stereotyped radial migration (Fig.9). Dbx1-derived glia is composed by GM protoplasmic, WM fibrous and WM radial astrocytes and distinctively express Reelin and Kcnmb4. While Dbx1 plays a fundamental role in V0 neuron identity (Pierani et al., 2001; Lanuza et al., 2004; Zagoraiou et al., 2009), here we demonstrate that it also controls astrocyte number through Notch signaling modulation.

**Figure 9.**
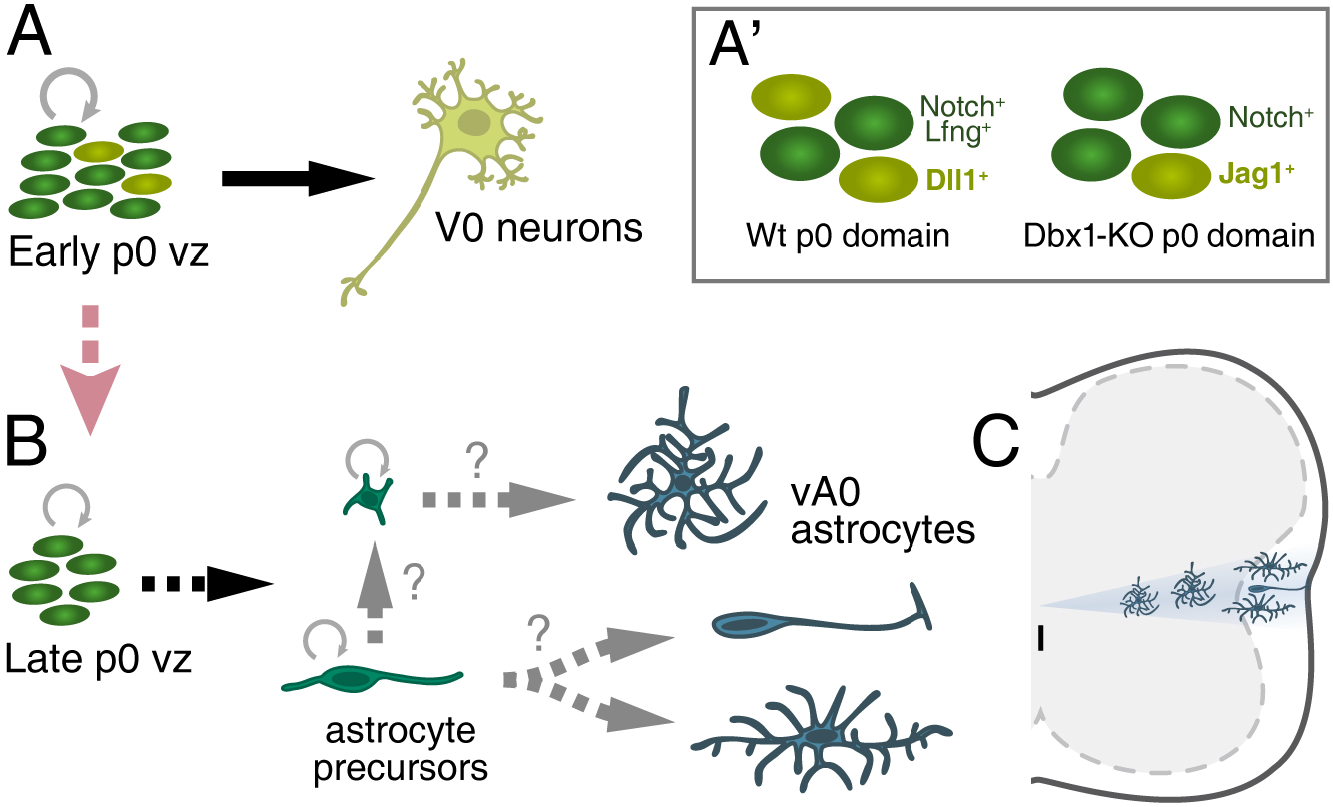
Dbx1 progenitors produce a heterogeneous population of spinal astrocytes. (A) Schematic representation of E10-E12 p0 domain and V0 interneurons. A’) In Dbx1-KO, neuronal precursors express Jag1 instead of Dll1, resulting in the expansion of the astrocyte progenitor pool. (B,C) During the gliogenic phase, the late p0 domain produces astrocytic precursors that migrate radially to colonize a specific region of the postnatal spinal cord. vA0 population comprises GM protoplasmic, WM fibrous and WM radial astrocytes.

### The precise allocation of vA0 cells and the formation of astroglial maps

We demonstrate that astrocytic cells derived from late p0 progenitors display an invariant positional arrangement in the mouse spinal cord. Mirroring the DV ventricular origin, vA0 population precisely occupies the entire intermediate portion of the spinal cord from the midline to the lateral funiculus, spatial restriction that is retained postnatally. Together with previous studies, the results presented here emphasize positional identity as a central developmental principle in the production of diverse glial populations of the spinal cord (Hochstim et al., 2008; Tsai et al., 2012; Vue et al., 2014). Like neuronal subtype specification, astrocyte diversification is highly influenced by the transcriptional code patterning DV progenitors (Hochstim et al., 2008; Muroyama et al., 2005; Vue et al., 2014; Zhao et al., 2014).

Interestingly, in line with descriptions in the mouse, a recent analysis of the midgestation human spinal cord have shown astrocytes with specialized DV transcriptional programs and gene expression signatures map also onto distinct anatomical domains (Andersen et al., 2021). In addition, studies in the Drosophila larval ventral nerve cord have shown that individual astrocytes are allocated to consistent positions with their arbors covering stereotyped territories of the neuropil (Peco et al., 2016). This developmental phenomenon conserved in humans, mice and flies suggests that astrocyte origin and final location is not incidental, and that spatially restricted groups of astrocytes are fitted to play specific roles in local neuronal circuits and microcircuits. In support of the connection between allocation and function, ventral horn astrocytes have been shown to play dedicated functions preserving motoneuron and sensorimotor circuit integrity (Molofsky et al., 2014; Kelley et al., 2018).

We found that Reelin and Kcnmb4 are novel WM vA0 molecular markers. Reelin was previously proposed to identify vA1 and vA2 WM astrocytes (Hochstim et al., 2008; Zhao et al., 2014). Here we show that Reelin is also present in Dbx1-derived astrocytes appearing after vA0s left the germinal zone. Its induction is likely dependent on Pax6, which persists in vA0 precursors and has the capacity to promote Reelin expression (Hochstim et al., 2008). Otherwise, Kcnmb4, encoding the β4 subunit of large conductance Ca^2+^-and voltage-activated K^+^ channels (BK), is already present in p0 ventricular precursors before their migration. It is intriguing to know the role of Kcnmb4 in WM vA0 astrocytes. In the brain, BK channels are located at astrocyte endfeet wrapping parenchymal vessels and pial arterioles (Contet et al., 2016; Filosa et al., 2006; Girouard et al., 2010). These channels, whose properties are adjusted by the β4 auxiliary subunit, are activated by neuronal stimulation to modulate smooth muscle contraction-relaxation, linking neuronal activity with local blood flow.

Our experiments show that the late Dbx1-progeny is exclusively composed of astroglial cells. This observation clashes with a previous study identifying oligodendrocytes being derived from DV intermediate vz region (Fogarty et al., 2005). This contradiction might result from the mapping strategies used, since in *Dbx1^lacZ^* mice, β-gal restricts to p0 cells, while the *Dbx1:Cre* transgene expression includes the dp5-dp6 domains (Fogarty et al., 2005), which are source of dorsal spinal oligodendrocytes (Cai et al., 2005; Vallstedt et al., 2005).

### vA0 astrocytes are morphologically heterogeneous

Sparse labeling of Dbx1-derived glia at perinatal stages revealed their morphological heterogeneity. vA0 population includes cells with features of the two classic astrocytic subtypes, protoplasmic and fibrous (Oberheim et al., 2012; Peters et al., 1991,Tabata, 2015 #225) (Fig.9B,C). Protoplasmic astrocytes have complex structures with numerous and highly branched processes and are intermingled with neurons in the intermediate spinal GM. Instead, vA0 cells with fibrous-like appearance are less complex with straight processes oriented longitudinally with axon fibers in the WM and express high levels of GFAP. The third vA0 morphological subtype is composed of radially oriented cells in the WM, also expressing GFAP and displaying conspicuous processes contacting the pia (Fig.9B,C). Cells with these characteristics have been recognized as important participants in the formation and maintenance of the glia limitans that covers the spinal cord surface (Liuzzi and Miller, 1987; Liu et al., 2013). In the adult, these radially-arrayed WM cells at the pial boundary have been shown to activate after autoimmune demyelination or contusive spinal cord injury (Petit et al., 2011).

Morphological astrocyte diversity appears to be a general characteristic of regionally-related astrocytic progenies. Several studies have shown that other astrocyte groups dorsal and ventral to vA0 also contain both GM and WM astrocytes (Tsai et al., 2012; Hochstim et al., 2008; Vue et al., 2014; Fogarty et al., 2005; Zhao et al., 2014). It remains to be established how GM and WM astrocytes are specified. Interestingly, in the dorsal spinal cord, Ascl1-expressing glial progenitors are restricted to become either GM or WM astrocytes or oligodendrocytes (Vue et al., 2014). Similarly, in the developing cortex, different astrocytic classes appear to emerge from separate clones (Garcia-Marques and Lopez-Mascaraque, 2013; Shen et al., 2021). Conversely, other clonal analyses also in the cortex have shown extensive morphological and location variability in astrocytes within clones (Clavreul et al., 2019), suggesting that cues from neighboring neurons influence astrocyte specialization (Farmer et al., 2016). Further studies will be required to determine whether individual p0 radial glial cells produce different astrocytic subtypes by symmetric or asymmetric division and how intrinsic and extrinsic cues act to sculpt vA0 astrocyte heterogeneity.

### Radial migration and proliferation in the parenchyma delineate vA0 spatial distribution

Dbx1-derived astrocyte precursors begin exiting the p0 vz domain around E13.5, and their number progressively rises until birth. Our experiments indicate that vA0 population depends primarily on cells emanating from the vz, and later on their proliferation in the parenchyma (Fig.9B). From E15.5 to E18.5, when the vz contribution is minor, vA0 population expands 3-4-fold. Such intense proliferation outside the germinal zone is a defining characteristic of astrocytic development, also shown in the cerebral cortex and in other regions of the spinal cord (Barry and McDermott, 2005; Tien et al., 2012; McMahon and McDermott, 2001; Ge et al., 2012). BrdU labeling shows that proliferation mainly takes place at distal-lateral positions, in the dGM and WM. Because more lateral territories covered by vA0s are larger than regions closer to the midline, they presumably require more cells. This distal expansion probably rely on the BRAF-MAPK pathway activated by local mitogens released from the basal lamina or secreted by neighboring neurons (Tien et al., 2012).

Direct observation of brain and spinal cord acute slices has shown that cells with a leading process attached to the pia travel through the tissue by continuous shortenings of its cellular extension (Nadarajah et al., 2001; Pakan and McDermott, 2014). Radial glial progenitors morphology with long cellular extensions spanning the thickness of the spinal cord, from their soma in the vz to the basal lamina, allows to anticipate vA0 migratory route toward the pia (McMahon and McDermott, 2002; Barry and McDermott, 2005). Dbx1-derived astroglial precursors progressively colonized lateral areas of the spinal cord following strictly radial movements. In this path from the vz to the parenchyma, we found that ~75% of vA0 precursors in the mantle are highly polarized and possess a prolonged extension toward the pia. This cell morphology and the gradual lateral displacement of nuclei suggest vA0 precursors mainly migrate translocating their soma. Further studies are needed to establish whether multipolar precursors, which account for ~25% of cells at E15-E16, are directly produced from the vz progenitors or if they arise through transformation of uni/bipolar precursors sowing the entire medial-lateral axis with vA0 cells (Fig.9B).

### Dbx1 regulates Notch ligand expression and neuron-astrocyte fate balance

The Notch signaling pathway plays a fundamental role in neuronal and glial development and undifferentiated pool preservation (Louvi and Artavanis-Tsakonas, 2006). High activation of Notch receptors in neural progenitors limits their neurogenic differentiation, while increasing glial commitment (Henrique et al., 1995; Chitnis, 1995; Gaiano et al., 2000; Park and Appel, 2003; Kong et al., 2015). Conversely, loss of key factors, such as Notch receptors or Rbpj triggers early increased neurogenesis and progenitor depletion (de la Pompa et al., 1997 231; Lutolf et al., 2002 285; Taylor et al., 2007; Kong et al., 2015). In addition, Notch signaling is known to regulate the transcription factors Nfia and Sox9, which are key players in glial commitment and gliogenesis onset (Deneen et al., 2006; Stolt et al., 2003; Kang et al., 2012). Notch activation in neural progenitors induces Nfia (Namihira et al., 2009) and collaborate on the maintenance of Sox9 expression (Taylor et al., 2007).

In the neural tube, Notch ligands Dll1 and Jag1 are expressed in discrete complementary DV territories, which define different quantitative levels of Notch signaling and neuron production rates in specific domains (Lindsell et al., 1996; Myat et al., 1996; Marklund et al., 2010; Ramos et al., 2010). We show that in the absence of Dbx1, the p0 domain switches from Dll1 to Jag1 expression (Fig.9A’), which is in line with patterning genes controlling the DV Notch ligand distribution (Skaggs et al., 2011; Marklund et al., 2010). In addition, *Dbx1-KO* progenitors lack the glycosyltranserase Lfng (Fig.9A’), which is known to suppress the ability of Jag1 to activate the pathway (Hicks et al., 2000; Yang et al., 2005; Kakuda and Haltiwanger, 2017). Thus, combined changes in both Notch-responding cells (Lfng^+^ to Lfng^−^) and ligand-presenting precursors (Dll1^+^ to Jag1^+^) contribute to enhance Notch signaling within the *Dbx1* mutant p0 domain, attenuating neuron differentiation and increasing glial production.

Our results highlight a role for Dbx1 in adjusting the size of p0-derived neuronal and glial populations. Strict control of neurogenesis is necessary not only to generate the appropriate number of V0 neurons, but also to preserve undifferentiated progenitors and their subsequent differentiation into vA0 cells. Dbx1-dependent expression of key Notch factors, together with the phenotypes in *Dbx1* mutants and after Notch manipulations, and the temporal coincidence of Dbx1 and Notch critical actions strongly support a mechanistic connection to control neuron-astrocyte cell fate balance.

## Materials and Methods

### Animals

All experiments involving animals were conducted according to the protocols approved by the Institutional Animal Care and Use Committee (IACUC) of the Fundación Instituto Leloir. Genotyping of *Dbx1*^*lacZ*^ (Pierani et al., 2001), *Glast*^*CreER*^ (Mori et al., 2006), *Nestin:CreER* (Carlen et al., 2006), *Psen1* (Shen et al., 1997), and *Ai14 tdTomato* conditional reporter (Madisen et al., 2010) mice were performed by PCR using allele-specific primers for each strain.

Time pregnancies were determined by detection of vaginal plug and midday was designated embryonic day (E) 0.5. Maximum induction of Cre activity in *Glast*^*CreER*^ mice was achieved with 150 mg/kg b.w. of tamoxifen (Tam, i.p. in corn oil, Sigma-Aldrich) administered to pregnant females. Mosaic labeling in transgenic *Nestin:CreER* mice was performed with Tam at a dose of 37.5 mg/kg b.w inyected at E11.75.

Embryos were dissected in PBS buffer. After decapitation, embryos were pinned on Sylgard plates, eviscerated and fixed for 1 h in 4% paraformaldehyde (PFA in PBS). They were cryoprotected in 20% sucrose (overnight, 4°C) prior to embedding in Cryoplast (Biopack). Stage-matched littermates of desired genotypes were aligned and embedded together to ensure identical processing conditions. Tissue was cryosectioned 30 μm, except in mosaic labeling experiments (45 μm) (Leica 3050S, Leica Biosystems).

For bromodeoxyuridine labeling, pregnant females or pups were injected with a single dose of BrdU i.p. 50 mg/kg b.w. and tissue was collected 3 hours later. Ly411575 γ-secretase inhibitor (Hyde et al., 2006) was administered to timed-pregnant dams at 5 mg/kg b.w. s.c (E10.5, E13.5 and E15.5 stages). Animals treated with vehicle (dimethyl sulfoxide in corn oil) served as controls in these experiments. The effectiveness of Ly411575 was controlled by appearance of tail shortening (when applied at E10.5) or haemorrhages in limbs and head (when injected at E13.5 or E15.5).

### Immunohistochemistry and in situ hybridization

Antibody stainings were performed essentially as previously described (Di Bella et al., 2019). Briefly, sections were treated with blocking solution (5% HI-serum, 0.1% TritonX-100 in PBS) for 1 h and primary antibodies in blocking were incubated overnight at 4°C. Antibodies used were: anti-Nkx2.2 (74.5A5, Developmental Studies Hybridoma Bank (DSHB), University of Iowa, USA; 1:20), anti-Nkx6.1 (F55A10, DSHB; 1:20), anti-Pax6 (cat.#pax6, DSHB; 1:20), anti-β-gal (55976, Cappel, now MP biomedicals, 1:1000 and from Martyn Goulding, Salk Institute; 1:1000), anti-Sox2 (SC17320, Santa Cruz Biotechnology; 1:300; MAB2018, R&D Systems 1:50), anti-Sox9 (AF3075, R&D Systems; 1:200), anti-Nfia (39397, Active Motif; 1:1000), anti-NeuN (A60, Chemicon; 1:500), anti-Olig2 (AB9610, Chemicon; 1:1000), anti-Gfap (from Fred Gage, Salk Institute, CA, USA; 1:2500), anti-Evx1 and anti-Pax3 (from Martyn Goulding, Salk Institute; 1:1000 and 1:400), anti-Nestin (from Fred Gage, Salk Institute; 1:500), anti-BrdU (MCA2060T, ImmunoDirect; 1:250), anti-Neurog2 (Sc-19233, Santa Cruz; 1:500), anti-Reelin (ab78540, Abcam; 1:500) and anti-dsRed (632496, Clontech; 1:300).

For detection, Cy-labeled species-specific secondary antibodies (Jackson ImmunoResearch) were incubated for 3-4 h at room temperature. For BrdU staining, acidic antigen retrieval was performed prior to immunodetection. Sections were mounted with PVA-DABCO or dehydrated in ethanol/xylene series and mounted with DPX.

Non-radioactive *in situ* hybridizations were performed as previously described (Carcagno et al., 2014). Sections were fixed with 4% PFA and washed with PBS-DEPC. Tissue was treated with proteinase K (3 μg/ml, 3 min), followed by 4% PFA for 10 min and PBS washes. Slides were incubated in triethanolamine-acetic anhydrate, pH 8.0, for 10 min and permeabilized with 1% TritonX-100 for 30 min. Sections were incubated for 2 h with hybridization solution (50% formamide, 5x SSC, 5x Denhardt’s solution, 250 μg/ml yeast tRNA). Digoxigenin (DIG)-labeled RNA probes used were Dbx1 and Dll1 (from Martyn Goulding, Salk Institute, CA, USA), Notch1 and Notch2 (from Geraldine Weinmaster, UCLA, CA, USA), Jag1, Dll3, Lfng and Kcnmb4 (this study). Dig-labeled probes were diluted in hybridization solution, denatured and incubated for 14 h at 68°C. Slides were washed three times, 45 min, at 68°C with 1x SSC, 50% formamide. For detection, sections were blocked with 10% HI-serum for 2 h and incubated overnight at 4°C with alkaline phosphatase-labeled sheep anti-dig antibody (11093274910, Roche; 1:2500). After washing, enzymatic activity was detected with BCIP and NBT (0.15 mg/ml and 0.18 mg/ml, Roche) in reaction solution (Tris pH 9.5 0.1M, MgCl_2_ 50mM, NaCl 0.1M, Tween-20 0.1%). For immuno-*in situ* double staining, sections were incubated with antibodies after developing the DIG *in situ* reaction.

β-gal activity was developed with X-gal using standard technique.

Images were captured by digital camera on Zeiss Axioplan microscope for brightfield or using Zeiss LSM5 Pascal and Zeiss LSM 510 Meta confocal microscopes and assembled using Zeiss ZEN, Adobe Photoshop and Adobe Illustrator.

### Quantitations and Statistical analysis

Quantification and analysis were performed on thoracic and upper lumbar spinal cord segments. Cell numbers are expressed per hemisection. The number of sections and embryos analyzed are indicated in the corresponding figure legend. For cell density maps and polar graphs, photos were aligned, the scatter plots with the position of cells were recorded with ImageJ and heat maps were constructed using a Matlab script (The MathWorks Co.). Differences between groups were evaluated by non-parametric Mann-Whitney U test or Kruskal-Wallis analysis of variance with *post hoc* Dunn’s Multiple Comparison test (GraphPad Software Inc). Results are presented as mean±SD and considered statistically significant when p<0.05.

## Acknowledgments

We thank Tom Jessell, Martyn Goulding, Jonas Frisén, Magdalena Gotz for mice; Flavio de Souza, Martyn Goulding, Pablo Vazquez, Alejandro Schinder and Fernando Pitossi for antibodies or probes, and Miguel Maroto for sharing Ly-411575. We deeply thank Abel Carcagno and Daniela Di Bella for insights and kind collaboration, and members of Schinder lab for discussions.

## Author Contributions

MMS performed most of experiments and analyzed the data; CAC collaborated with some experiments; GML designed the research and performed some experiments. MMS and GML prepared the figures and wrote the manuscript.

## Funding

This work was supported by the Agencia Nacional de Promoción Científica y Tecnológica of Argentina (PICT2014-1821 and PICT2017-0297) to GML.

## Competing interests statement

The authors declare no competing financial interests.

## Notes

### Competing Interest Statement

The authors have declared no competing interest.

